# Structure and function of a cross-neutralizing influenza neuraminidase antibody that accommodates recent N2 NA Asn245 glycosylation

**DOI:** 10.1101/2025.06.30.662356

**Authors:** Xueyong Zhu, Ahmed M. Khalil, Michael S. Piepenbrink, Wenli Yu, Yao Ma, Luis Martinez-Sobrido, Ian A. Wilson, James J. Kobie

**Author notes:** contributed equally to the work (Xueyong Zhu, Ahmed M. Khalil, Michael S. Piepenbrink). Present Address: Department of Biomedical Sciences, The College of Medicine, Florida State University, Tallahassee, FL 323094, USA.

## Abstract

Monoclonal antibodies (mAbs) that recognize and inhibit a diverse range of influenza viruses, although relatively rare, have been isolated following infection or vaccination. Study of their ontology and mechanisms of action informs universal vaccine and therapeutic development. We have previously described a potent and broad neuraminidase (NA)-neutralizing human mAb, 1122A11, that neutralizes a wide range of H3N2 viruses. Here, further characterization of 1122A11 reveals reactivity to cross-group influenza A virus NAs, including group-1 N1 and N8, and group-2 N2 and N3 NAs. Recent H3N2 viruses have acquired Asn245 glycosylation on the active site rim. Crystal structures of an N2 NA from A/Singapore/INFIMH-16-0019/2016 (H3N2) at 2.3 Å (apo) and 2.2 Å (Fab bound) resolution showed that 1122A11 binding causes local changes to the periphery of NA active site to accommodate the glycan. The CDRH3 of 1122A11 inserts into the active site and mimics the substrate sialic acid. We then determined that the ability of 1122A11 to protect from lethal challenge in mice is not dependent on Fc-effector function. These results highlight the therapeutic potential of 1122A11 as a broad protective anti-viral and reinforce pursuit of immunogen development of NA antibodies toward achieving more universal influenza protection.

## Introduction

The antigenic drift and shift of influenza A viruses (IAV), and the tendency for natural infection and current vaccines to induce predominantly strain-specific antibodies, results in global susceptibility to new strains of seasonal IAV, and great risk from IAV with pandemic potential for which minimal or no immunity may exist. Generally, H3N2 IAV has undergone a higher rate of antigenic drift than H1N1 IAV and influenza B virus (IBV) strains^1, 2^ and, in recent years (2016-2022), incomplete match between vaccine and circulating strains and amino acid and glycosylation site changes in the hemagglutinin (HA) of egg-adapted strains likely contributed to the modest 5%-36% vaccine efficacy^3, 4, 5^. However, neuraminidase (NA) undergoes less antigenic drift compared to HA^2, 6^, and its enzymatic active site, which mediates hydrolysis of sialic acid to enable viral budding and progeny escape, is highly conserved. Thus, neuraminidase has emerged in recent years as an attractive target for vaccine and therapeutic development^7, 8, 9^.

In the 2014-2015 influenza season, a novel Asn245 glycosylation of N2 NA was first detected in human H3N2 viruses and, since 2016-2017 season, almost all circulating seasonal human H3N2 virus NAs carried such glycosylation^10, 11^. This Asn245 glycan is located near the NA active site and blocks binding and inhibition of some NA active site-specific antibodies^10, 11, 12^. Although the inhibition ability was reduced, some NA active site antibodies such as broad IAV and IBV protective 1G01-like antibodies still maintain binding and protective efficacy^12^. Previously, low-resolution negative-stain electron microscopy has shown 1G01-like antibody binding to the N2 NA with Asn245 glycosylation, although with reduced inhibitory activity^12^, but the detailed mechanism of how such antibodies overcome Asn245 glycosylation still need further investigation.

Antibodies can mediate anti-viral effects through various processes including those dependent on Fc-effector functions, by which the binding of antibody to a virus is recognized by Fc receptors expressed on innate cells (e.g. macrophages and NK cells) or by complement proteins, and subsequently mediate viral clearance. Additionally, some antibodies that recognize critical viral epitopes and mediate virus attachment or other critical processes can directly neutralize the virus without a requirement for accessory cells or proteins for their anti-viral activity. Several monoclonal antibodies (mAbs) have been reported that have broad activity against a wide range of influenza viruses, including those that target the stem region of HA, which appear to be at least partially dependent on Fc-mediated effector functions for its *in vivo* activity^13, 14, 15, 16^, and which have driven major efforts in developing vaccines that promote the development of HA stem-targeting antibodies^17, 18^. In recent years, NA-specific mAbs that have broad antiviral activity have also been described with variable dependence on Fc-mediated effector functions, such as broad N2-specific NA-head underside antibodies 3C08, 3A10 and 1F04 all confer protection with Fc effector function^19^, while the IAV and IBV NA active site antibody FNI9 does not appear to be dependent on Fc effector function^20^.

We have previously described 1122A11, an NA-specific human mAb isolated from peripheral blood plasmablasts from a participant vaccinated with the 2014-2015 seasonal quadrivalent vaccine based on influenza viral strains A/California/07/2009 (H1N1), A/Texas/50/2012 (H3N2), B/Massachusetts/2/2012, and B/Brisbane/60/2008 ^21^. 1122A11 potently neutralizes a wide range of H3N2 viruses, inhibits their NA enzymatic activity, and has prophylactic and therapeutic activity against lethal challenge in mice^21^. Here, we further characterize 1122A11 to be a cross-group anti-NA antibody. To determine the structural and functional basis for its anti-viral activity, the crystal structure of 1122A11 Fab in complex with the N2 NA from A/Singapore/INFIMH-16-0019/2016 (H3N2) (Singapore16 N2), which contains the recent Asn245 glycosylation site, revealed that 1122A11 binds to the NA active site, mimicking in part the substrate sialic acid. Functional studies indicated that the antibody *in vivo* activity is not dependent on Fc-effector function.

## Results

### mAb 1122A11 exhibits cross NA group binding and inhibition

Antibody 1122A11 was previously characterized to be an anti-NA antibody against a wide range of N2 NAs. Here, we sought to determine the breadth of its recognition and inhibition of NAs from IAV and IBV viruses. In an enzyme-linked immunosorbent assay (ELISA) and NA enzymatic inhibition assay using substrate 2′-(4-Methylumbelliferyl)-α-D-*N*-acetylneuraminic acid (MUNANA), 1122A11 demonstrated binding to recombinant IAV group-1 N1 and N8, and group-2 N2 and N3 NAs (Fig. 1 and Supplementary Fig. 3). Unlike the well described NA active site mAbs that broadly neutralize IAV and IBV NA subtypes, such as 1G01 and 1E01^7^ as well as FNI9^20^, 1122A11 had minimal or no detectable binding to tested IAV N4, N5, N6, N7 and IBV NAs (Fig. 1a). The breadth of 1122A11 to inhibit NA enzymatic activity using the small substrate MUNANA (Fig. 1b) largely mirrored its binding profile with corresponding NAs tested.

**Fig. 1.**
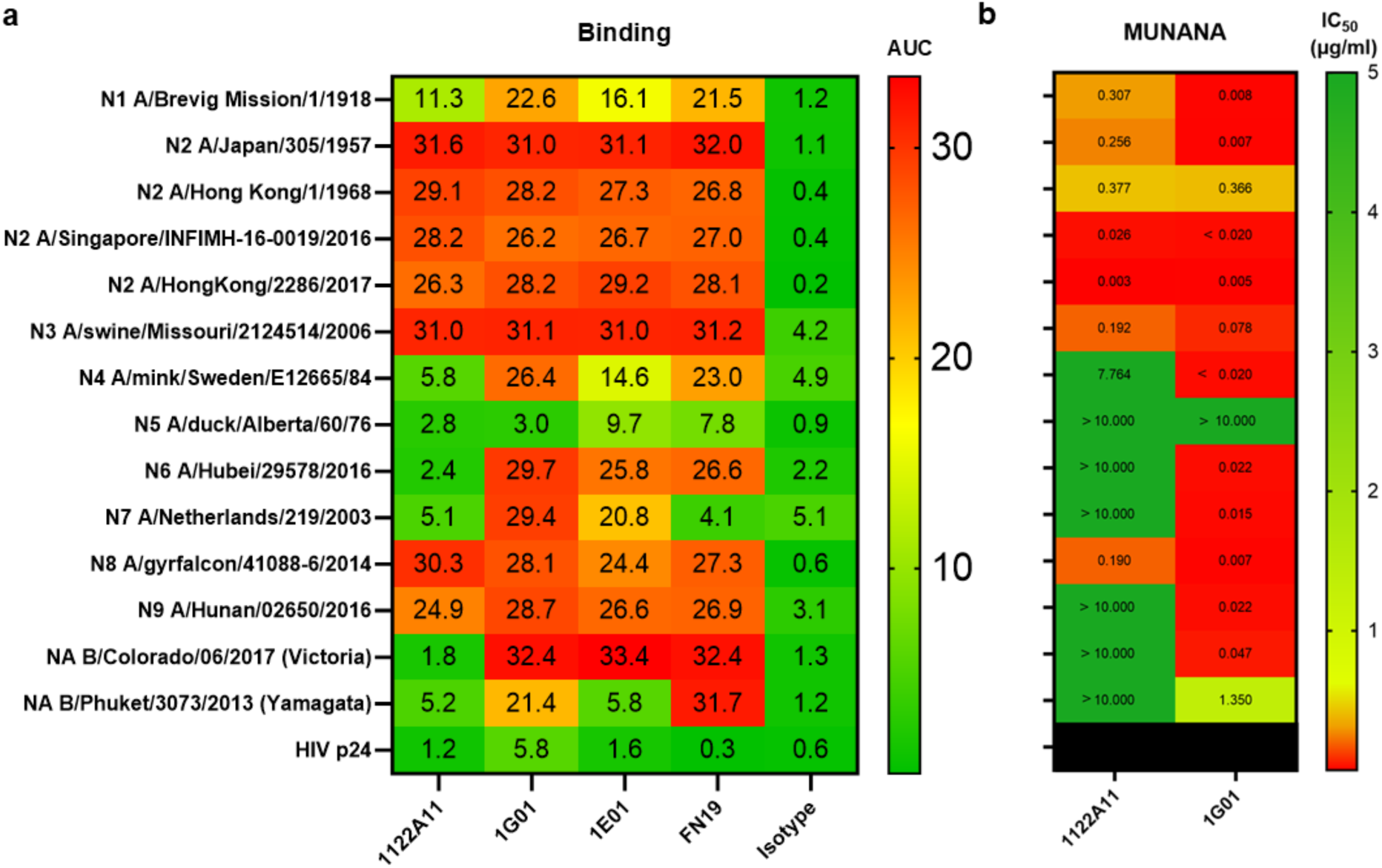
Breadth of mAb 1122A11. **a** Heat map of 1122A11 binding (presented as area under the curve (AUC) with red representing highest binding and green lowest binding across all data) to recombinant NA proteins in the presence of calcium and magnesium in ELISA with positive control pan-NA antibodies 1G01, 1E01 and FNI9, as well as a negative control antibody Isotype. **b** Heat map of antibody inhibition (with red representing lowest IC_50_ concentration and green the highest IC_50_) to the same recombinant NA proteins in MUNANA assay with positive control antibody 1G01.

### Overall structure of 1122A11 Fab in complex with Singapore16 N2 NA

To elucidate how 1122A11 broadly protects against influenza virus infection, we determined crystal structures of 1122A11 Fab in complex with Singapore16 N2 NA ectodomain (residues 82-469) at 2.2 Å resolution (Fig. 2, Supplementary Table 1) and apo Singapore16 N2 NA at 2.3 Å resolution (Fig. 3, Supplementary Table 1). Singapore16 H3N2 virus was the vaccine strain for the 2018-2019 influenza season in which the N2 NA introduced an *N*-linked glycosylation site at residue Asn245 contributing to antigenic drift^10^ and blocking or reducing inhibition by antibodies^11, 12^.

**Fig. 2.**
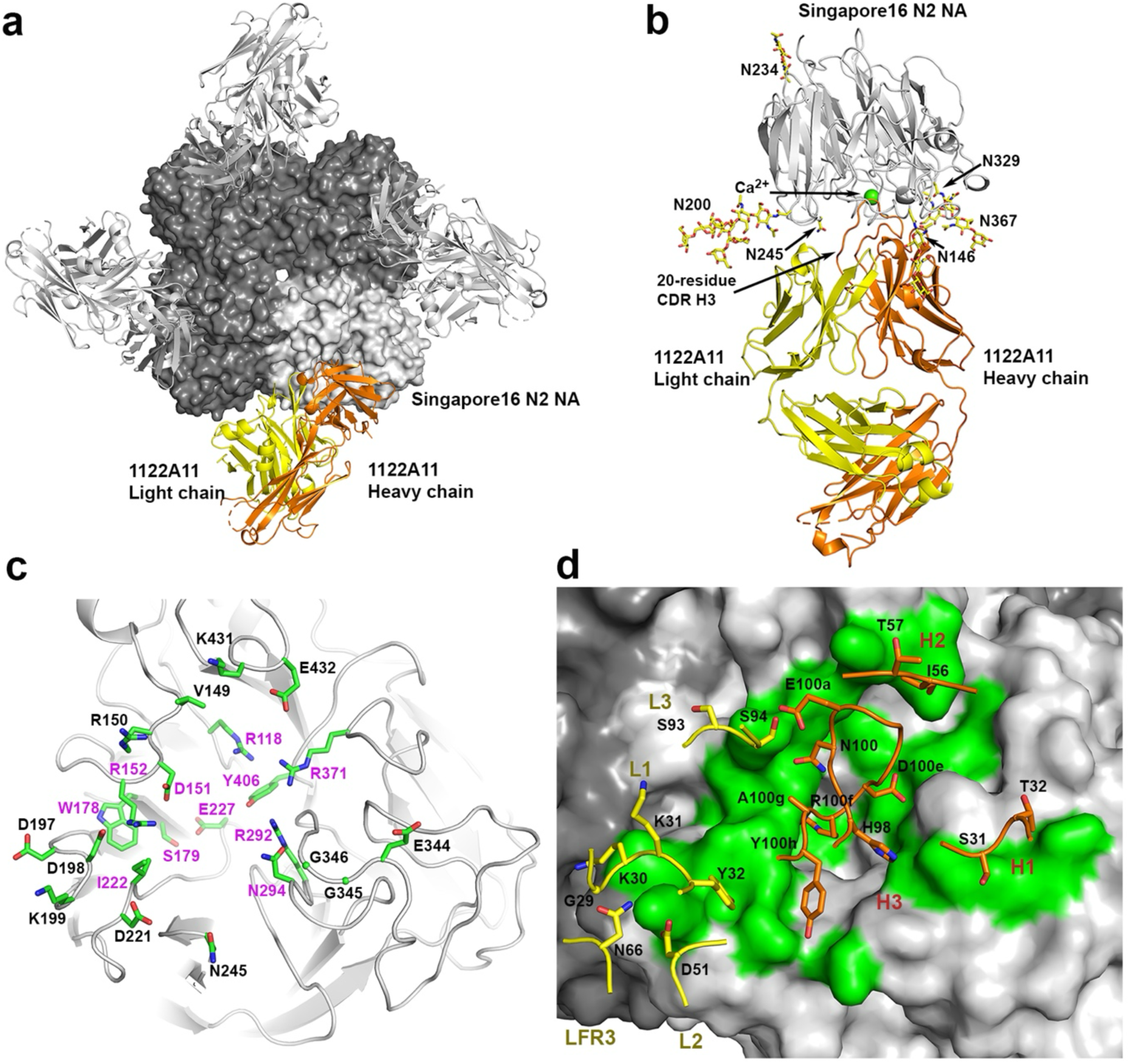
Crystal structure of 1122A11 Fab in complex with Singapore16 N2 NA at 2.2 Å resolution. **a** The NA tetramer with one Fab bound to each NA protomer. **b** An NA protomer bound to one Fab. **c** The antibody epitope residues on the NA. **d** The antibody paratope residues superimposed on the NA surface in a similar view to panel (**c**). For all panels, one NA-Fab protomer is colored with NA in light grey, and Fab light chain in yellow, and Fab heavy chain in orange, and the NA and Fab in other NA protomers are in dark and light grey, respectively. *N*-linked glycans in (**b**) are represented in sticks with yellow carbon atoms. A bound calcium ion in (**b**) is shown as a green sphere. The epitope residues labeled in pink in (**c**) are conserved residues in influenza A N1-N9 and influenza B NA subtypes. The molecular surface depicting the epitope in (**d**) is colored in green.

**Fig. 3.**
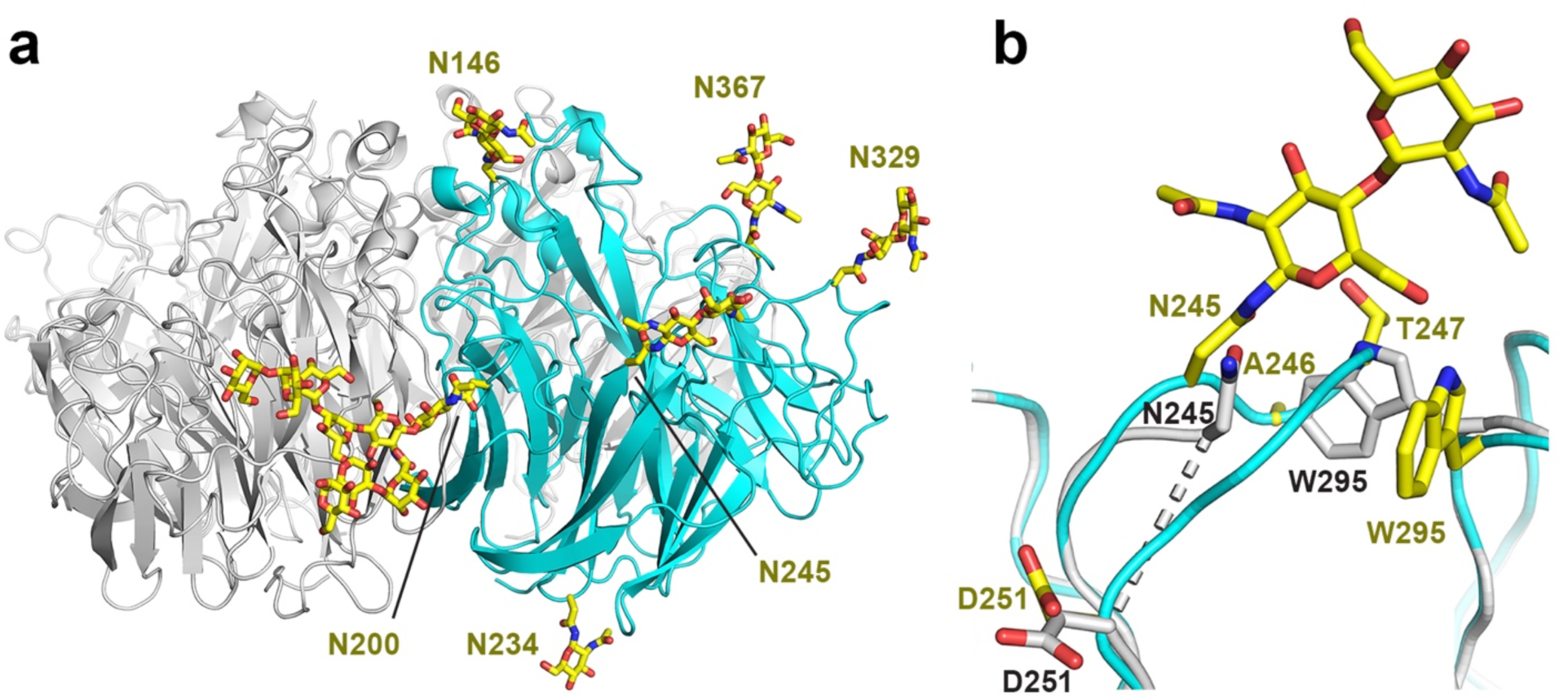
Crystal structure of apo form Singapore16 N2 NA at 2.3 Å resolution. **a** The NA tetramer with one protomer in cyan with *N*-linked glycans represented in sticks with yellow carbon atoms. **b** Structural comparison of the apo form (unliganded) of Singapore16 N2 NA (in cyan and yellow-carbon sidechains and glycans) with its antibody-bound form (in grey) around the Asn245 glycosylation site.

Antibody 1122A11 binds Singapore16 N2 NA tetramer with one Fab per NA protomer via both heavy and light chains (Fig. 2a, b), burying 660 Å^2^ of NA surface where 71% arises from the heavy chain and 29% from the light chain. Twenty-three residues in N2 NA active site (Fig. 2c) interact with all heavy chain complementarity determining regions (CDRs) H1, H2, H3 and light chain CDRs L1, L2 and L3, as well as light chain framework region 3 (LFR3) (Fig. 2d). However, CDR H3 dominates the Fab-NA interactions (Fig. 2d).

### Singapore16 NA epitope residues for NA specificity of 1122A11

Antibody 1122A11 blocks the N2 NA active site (Fig. 2), which is consistent with its inhibition of NA enzymatic activity in both ELLA and NA-Star experiments reported previously^21^ as well as in the NA MUNANA-based assays in this study (Fig. 1b). Around half of the Singapore16 N2 NA residues within the 1122A11 epitope are highly conserved among all IAV N1-N9 and IBV NA subtypes (Fig. 2c, Supplementary Table 2), including seven of eight charged and polar catalytic residues in the active site, Arg118, Asp151, Arg152, Arg292, Asn294, Arg371 and Tyr406 (except Glu276, which is not an epitope residue), as well as four conserved framework residues for stabilizing the active site, Trp178, Ser179, Ile222 and Glu227^22, 23^. Moreover, framework residue Asp198 is also conserved in the IAV N1, N2, N3, N4, N5 and N8 as well as IBV NAs that we tested (Fig. 2c, Supplementary Table 2).

The 1122A11 20-residue CDR H3 mainly interacts with conserved residues in the active site, whereas CDRs L1, L2, L3, H1 and H2, as well as framework LFR3, largely contact non-active site residues (Figs. 2d and Supplementary Fig. 1). Among hydrogen bonds and salt bridges between the Fab and the NA, CDR H3 contributes most interactions with salt bridges between antibody and NA residues, including H3 Glu100a with NA Arg150, H3 Asp100e with NA Arg118, Arg292 and Arg371, and H3 Arg100f with NA Glu227 and Asp151. CDR H3 Asn100, Arg100f, Ala100g main-chain, and Tyr100h make also H-bonds with NA Asp151, Trp178 main-chain, Arg152, and Asp221, respectively (Supplementary Fig. 1). Among other key antibody-NA interactions, CDR L1 Gly29 and Lys30 as well as CDR L2 Asp51 make H-bonds and a salt bridge with NA Lys199, L1 Tyr31 forms a H-bond with NA Asp221, and H2 Thr57 form main-chain H-bonds with NA Lys431 (Supplementary Fig. 1).

### 1122A11 renders NA Asn245 glycosylation site disordered when it binds Singapore16 N2 NA

To compare with Singapore16 N2 NA in its complex with 1122A11 Fab, we determined crystal structures of the head domain of the apo (unliganded) N2 NA at 2.2 Å resolution (Fig. 3). The expected N-linked glycans of apo Singapore16 N2 NA all had interpretable electron density including Asn245 as well as Asn146, Asn200, Asn234, Asn239 and Asn367 (Fig. 3a). However, upon 1122A11 binding, the N-linked glycan at Asn245 could not be modeled at atomic resolution, although weak electron density for the glycan was present, whereas other potential N-glycans had interpretable electron density and could be modeled (Fig. 2b). Furthermore, the N2 NA in the Fab-NA complex structure was highly ordered except for loop residues 246-250 (Ala-Thr-Gly-Lys-Ala) following Asn245 that were disordered (Fig. 3b).

### Substrate mimicry by 1122A11

In 1122A11, CDR H3 appears to have the most important role in NA binding by inserting into the active site pocket (Fig. 2 and Supplementary Fig. 1). Among CDR H3 paratope residues, the adjacent Asp100e and Arg100f residues interact with positively and negatively charged catalytic residues (Fig. 4a). The Asp100e carboxylate closely aligns with the carboxylate of the substrate sialic acid, and engages a conserved critical trio of NA arginine residues, Arg118, Arg292 and Arg371, using a similar salt-bridge network (Fig. 4a, b). On the other hand, Arg100f forms salt-bridges with conserved NA Asp151 and Glu227, and H-bonds with the main-chain carbonyl of conserved Trp178 (Fig. 4a). NA substrate mimicry of antibody CDR H3 loops has been reported for other human NA active site mAbs. Cross-reactive anti-N1 and N9 NA active site mAb Z2B3^24^ uses two similar adjacent H3 residues, Asp100c and Arg100d. IBV NA active site mAb 1G05^25^ has H3 Asp100a and Arg100b that interact with the same conserved NA active-site residues using similar salt-bridge and H-bond networks as 1122A11 (Fig. 4c, d). Interestingly, pan-NA broad neutralizing mAb FNI9^20^ also harnesses two CDR H3 aspartic and arginine residues, Arg100b and Asp100c, but in opposite directions on H3, to form similar salt-bridge and H-bond networks as observed in 1122A11, Z2B3 and 1G05 (Fig. 4a). Anti-N9 active site mAb NA-45 was the first reported anti-NA antibody structure to show substrate mimicry^26^. Instead of using an aspartic residue, NA-45 H3 Glu100a forms a similar salt-bridge network with the same tri-arginine residues in the N9 NA active site, while H3 residue100b is a leucine instead of arginine and does not make the same interactions (Fig. 4f). All of these NA mAbs with substrate mimicry have relatively long CDR H3 loops (defined by the third residue following Cys92 to the residue preceding Trp103 in Kabat numbering) for 1122A11 (20 residues), Z2B3 (23 residues), 1G05 (17 residues), FNI9 (19 residues) and NA-45 (17 residues) that enable insertion deep into the active site.

**Fig. 4.**
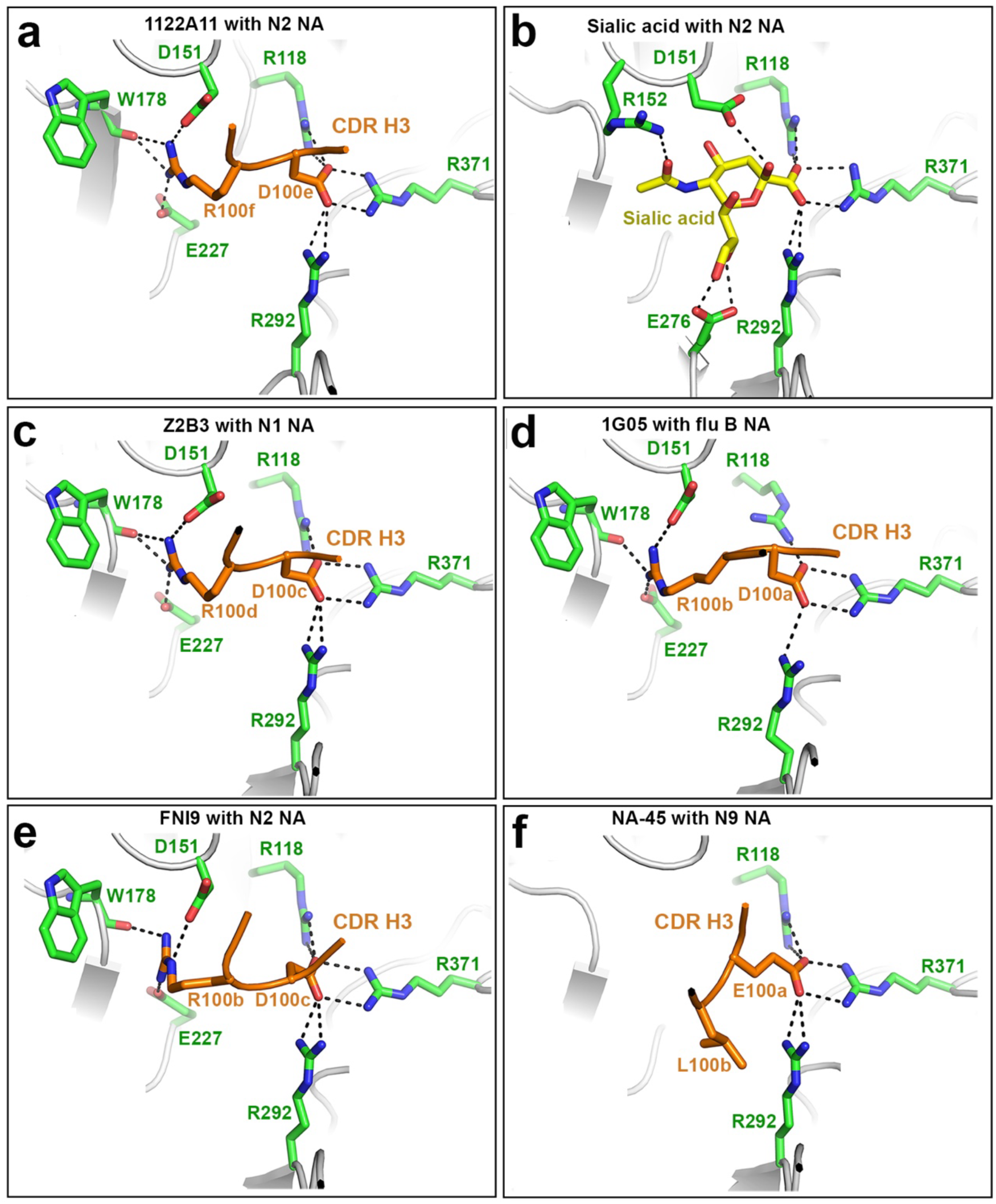
Comparison of NA active-site interactions with 1122A11 and other mAbs that demonstrate substrate mimicry. **a** 1122A11 H3 D100e and R100f with Singapore16 N2 NA. **b** Sialic acid with N2 NA from A/Tanzania/205/2010 (H3N2) (PDB code 4GZQ). **c** Z2B3 H3 D100c and R100d with N1 NA from A/Anhui/1/2013 (H7N9) (PDB code 6LXJ). **d** 1G05 H3 D100a and R100b with influenza B NA from B/Phuket/3073/2013 (PDB code 6V4N). **e** FNI9 H3 R100b and D100c with N2 NA from A/Tanzania/205/2010 (H3N2) (PDB code 8G3N). **f** NA-45 H3 E100a and L100b with N9 NA from A/Shanghai/2/2013 (H7N9) (PDB code 6PZE). For comparison, mAbs are numbered in the Kabat scheme, and NAs are numbered in N2 NA numbering. Accordingly, mAb Z2B3 and the N1 NA in PDB 6LXJ (**c**), influenza B NA in PDB 6V4N (**d**), and mAb FNI9 in PDB 8G3N (**e**) are renumbered.

### 1122A11-LALAPG mutant has comparable binding to N2 with 1122A11

After determining that 1122A11 does indeed directly bind to the NA active site, we sought to investigate if it can directly mediate *in vitro* and *in vivo* anti-viral activities without reliance on Fc-effector functions. Subsequently, the ability of 1122A11 IgG1 to bind to FcγRs was minimized by incorporating the well-described L234A and L235A substitutions (LALA) to residues in the C_H_2 domain, which form part of the FcγR binding site, and the P329A substitution (PG) in the C1q binding site to minimize binding to C1q^27, 28^. The resulting 1122A11-LALAPG, as expected, had minimal binding to the FcγRI as compared to 1122A11 (Fig. 5a). 1122A11-LALAPG retained its ability to bind to N2 NA from A/Hong Kong/2286/2017 (H3N2), with no significant differences compared to 1122A11 or 1G01^7^, another NA-specific mAb (Fig. 5b).

**Fig. 5.**
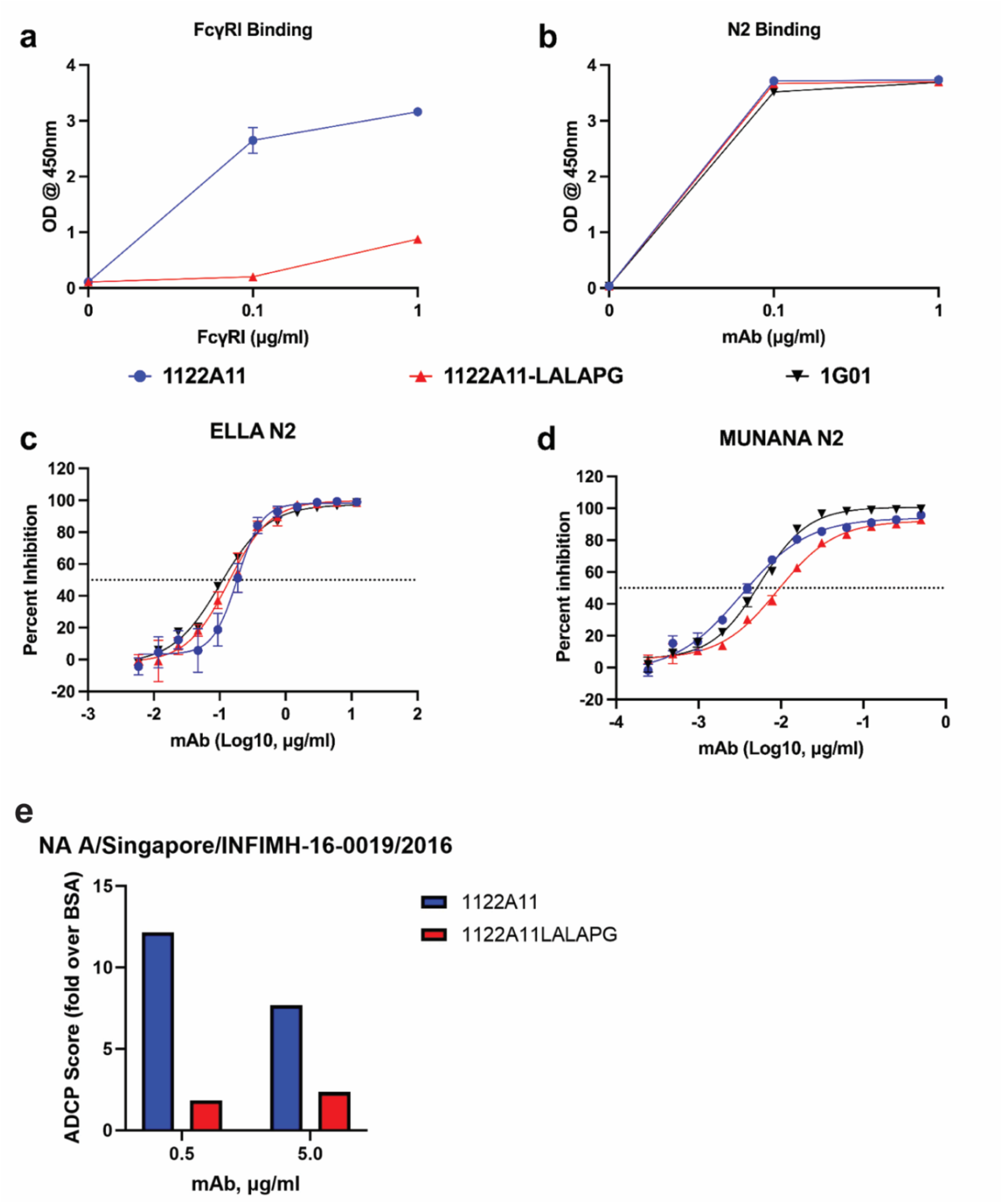
Functional profile of 1122A11-LALAPG. **a** 1122A11 and 1122A11-LALAPG were tested in triplicate at 1 μg/mL for binding to increasing concentrations of FcγRI by ELISA. **b** Increasing concentrations of 1122A11 and 1122A11-LALAPG were tested in triplicate for binding to recombinant NA proteins of A/Hong Kong/2286/2017 by ELISA. Inhibition of recombinant NA (A/Hong Kong/2286/2017) activity was compared between 1122A11 and 1122A11-LALAPG. The recombinant NA was added with serial dilutions of the mAbs, in duplicate, in ELLA (**c**) and MUNANA (**d**) assays. Triplicate and duplicate data are presented as mean ± SD (error bars). **e** Ab-dependent cellular phagocytosis (ADCP) assay. NA-coated and BSA-coated beads were incubated with hMAb and then added to THP-1 cells. After incubation, cells were assayed for fluorescent bead uptake by flow cytometry. The ADCP score of each hMAb was calculated by multiplying the percentage of bead-positive cells (frequency of phagocytosis) by the mean fluorescence intensity (MFI) of the beads (degree of phagocytosis) and dividing by 10^6^.

### 1122A11-LALAPG retains ability to inhibit NA enzymatic activity and neutralize H3N2 virus

The primary function of influenza NA is to cleave sialic acid residues from the cell surface, enabling release of progeny virus, which can then infect new cells. Consistent with our previous findings that 1122A11 can potently inhibit NA enzymatic activity^21^, 1122A11–LALAPG inhibited NA activity in the ELLA assay (Fig. 5c).1122A11 and 1122A11–LALAPG both potently inhibited the activity of recombinant NA, with comparable IC_50_ of 0.0184 and 0.0141 μg/mL, respectively. The inhibition of NA enzymatic activity was further confirmed in the small substrate MUNANA assay, with 1122A11 and 1122A11–LALAPG having comparable IC_50_ of 0.003 and 0.008 μg/mL, respectively (Fig. 5d). These results indicate that 1122A11 inhibition of NA enzymatic activity is not Fc-dependent. The effectiveness of the LALAPG mutations to reduce Fc-effector function was then tested in an antibody dependent cellular phagocytosis assay. As expected, the ADCP activity of 1122A11 LALAPG was greatly reduced compared to 1122A11 (Fig. 5e). The ability of 1122A11-LALAPG to inhibit viral infection was then measured using a microneutralization assay. Both 1122A11 and 1122A11-LALAPG were able to neutralize A/Uruguay/716/07 (H3N2) (Supplementary Table 3), with comparable NT_50_ of 1.56 and 0.78 μg/mL respectively, which were slightly lower (more potent) than the control 1G01 and DA03E17^29^ NA active site mAbs. These results indicate that *in vitro* neutralization of H3N2 virus by 1122A11 is not affected by these mutations in the Fc fragment. To further determine if the Fc effector function is necessary for *in vitro* neutralization, F(ab’)2 of 1122A11 was compared to intact 1122A11 and the LALAPG mutant. Comparable NT_50_ values were determined for all three (Supplementary Table 4) indicating that Fc effector function is not required for *in vitro* viral neutralization.

### 1122A11-LALAPG protects from lethal H3N2 infection

Determining that 1122A11 directly binds the active site of N2, potently inhibits its enzymatic activity, and neutralizes H3N2 viruses *in vitro*^21^, we sought to determine if its *in vivo* efficacy was dependent on Fc-effector function. Mice received a single intraperitoneal injection of 1122A11 or 1122A11-LALAPG prior to challenge with a lethal dose of H3N2 X31 virus. All H3N2 X31-infected control mice (isotype control and mock-treated groups) required euthanasia by 5 days post-infection (p.i.), while all 1122A11-and 1122A11–LALAPG-treated mice survived infection (Fig. 6a). All H3N2 X31-infected mice that were treated with isotype control human mAb or mock-treated had progressive weight loss that reached the 75% endpoint threshold by 5 days p.i. (Fig. 6b). Only minimal to modest weight loss occurred in mice treated with 1122A11 or 1122A11-LALAPG, with no significant differences between 1122A11 or 1122A11-LALAPG, although a trend toward greater weight loss with 1122A11-LALAPG compared to 1122A11 at similar doses. All doses of 1122A11 and 1122A11–LALAPG resulted in significant reduction in lung virus titers at 2 and 4 dpi, with no significant difference between equal doses of 1122A11 and 1122A11–LALAPG (Fig. 6c). Together, these results indicate that 1122A11 and 1122A11–LALAPG provide similar protection from lethal H3N2 infection, suggesting 1122A11 can mediate effective protection against H3N2 without dependence on Fc-effector functions.

**Fig. 6.**
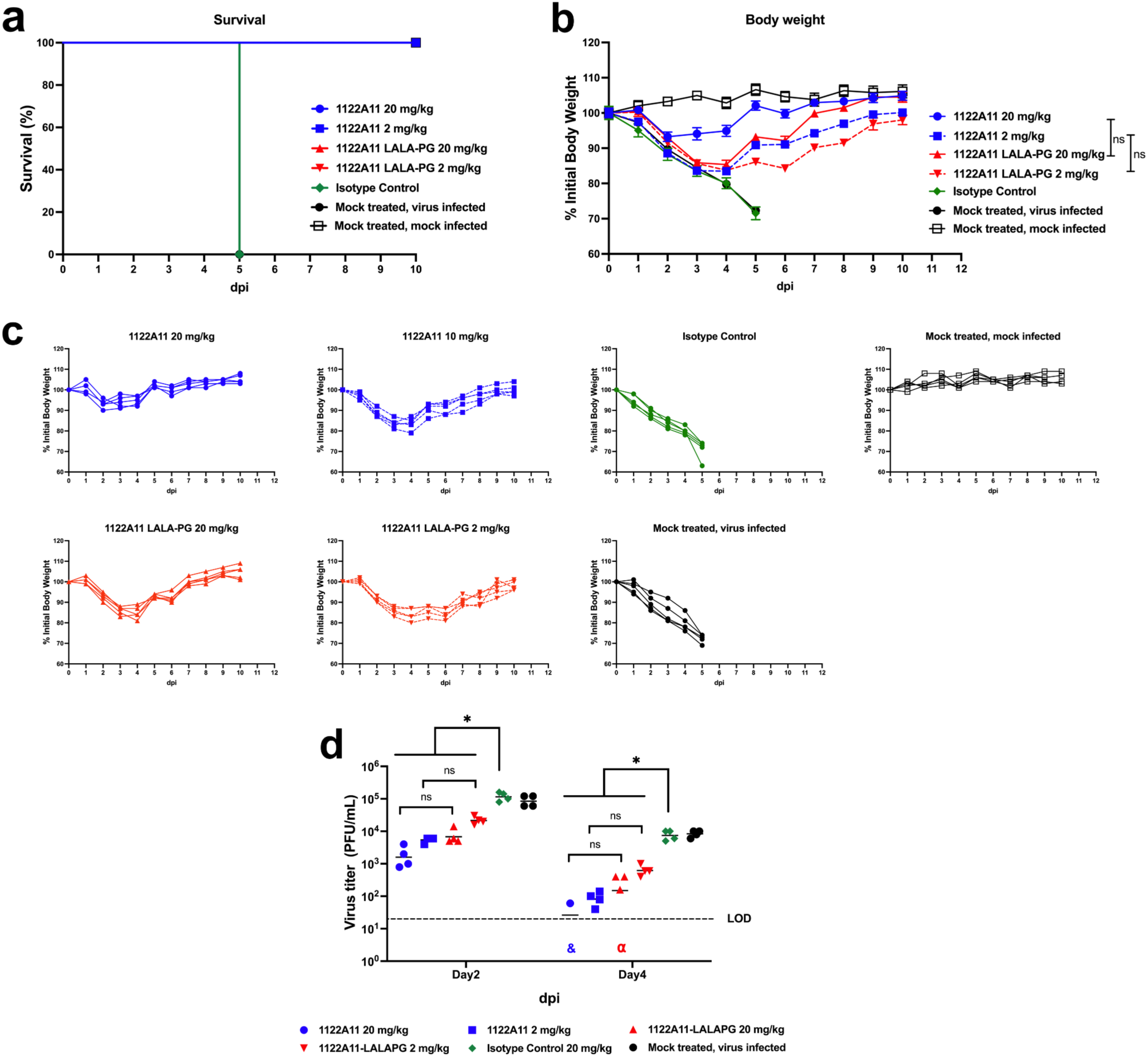
*In vivo* prophylactic activity of 1122A11-LALAPG. C57BL/6 mice (7-weeks-old) were injected with either 20 or 2 mg/kg of indicated mAbs or PBS (mock-treated), intraperitoneally, 24 h before infection. Mice were then challenged with 10X MLD_50_ (3×10^4^ PFU) of X31 and monitored daily for 10 days p.i. for survival (**a**) and body weight loss displayed in summary (mean ± SD) (**b**) or individual (**c**) (N = 5, group). Significance between equal mAb doses were determined via two-way ANOVA. To evaluate viral replication in the lungs (**d**), mice were sacrificed at 2 (N = 4, group) and 4 (N = 4, group) days p.i., and whole lungs were used to quantify viral titers by standard plaque assay (PFU/mL). Each symbol represents an individual mouse. * indicates significant differences (p < 0.05 using one-way ANOVA and Tukey’s test). “ns” indicates non-significant statistical difference (p > 0.05). Dotted line indicates the limit of detection (LOD) of the assay (20 PFU/mL). The “&” symbol indicates that the virus was not detected in three mice per group. The “α” symbol indicates that the virus was not detected in one mouse per group.

## Discussion

There is a pressing need for new influenza vaccines that can effectively withstand the ongoing mutations in circulating human viruses and confer universal protection. The promising outcomes observed in pre-clinical and clinical studies involving NA-directed antibodies and antivirals underscore the therapeutic promise associated with targeting NA. Here we determined that mAb 1122A11 isolated from a seasonal influenza vaccine participant is reactive to IAV group-1 N1 and N8 as well as group-2 N2 and N3 NAs, due to its binding to the highly conserved NA active sites. However, although NA active-site pocket residues are conserved across subtypes, the active site rim residues are more diversified (Supplementary Table 2), and in particular, residues surrounding the 1122A11 epitope, 344-346 and 431-432, exhibit insertions and deletions among different NA subtypes. The variability consequently restricts 1122A11 binding to other tested N4, N5, N6, N7, N9 and influenza B NAs. The active site rim residue variability also explains why neutralizing antibodies targeting the active site show varying breadth against different NA subtypes: they interact with distinct epitopes associated with these active site rim residues (Supplementary Fig. 2, Supplementary Table 2). For instance, 1122A11 targets to a range of IAV NAs, whereas mAbs such as 1G01^7^, 1E01^7^ and FNI9^20^ are reactive across IAV and IBV NA subtypes. Further studies, including mutagenesis analyses, are required to precisely identify the NA residues responsible for 1122A11’s limited breadth.

NA sialic acid substrate mimicry appear to be important for anti-NA antibody breath and potency, as the key interacting NA active-site residues are evolutionary conserved for NA enzymatic activity. Several anti-NA active site antibodies such as 1122A11, Z2B3, 1G05, FNI9 and NA-45 use substrate mimicry by forming salt-bridges with three conserved arginine residues, Arg118, Arg292 and Arg371 using Asp or Glu residues in CDR H3 to mimic the sialic acid carboxyl group.

For NAs of circulating influenza H3N2 viruses, in years following the 2014-2015 season, almost all N2 NAs in humans carry an N-glycosylation site at Asn245, which is confirmed in our crystal structure of apo Singapore16 N2 NA. The Asn245 glycosylation of N2 NA has been shown to block binding of a number of antibodies that target the NA active site^10, 11^. Nevertheless, although 1122A11 was discovered from a vaccinee immunized with the 2014-2015 seasonal quadrivalent vaccine which included viral strain A/Texas/50/2012 (H3N2) that did not possess N2 NA site 245 N-glycosylation, it is still reactive to N2 NAs with Asn245 N-glycosylation including Singapore16 N2 NA akin to 1G01 and 1G01-like antibodies^12^. It was previously determined that 1G01 still binds to N2 NA active site by low-resolution negative stain electron microscopy but with lower inhibitory activity^12^. Here, from the high-resolution crystal structure of 1122A11 with Singapore N2 NA, we determined that 1122A11 can bind to the NA in the presence of Asn245 glycosylation through promoting disorder of the glycan moiety of glycosylated Asn245 and the following NA loop residues 246-250 compared to apo Singapore N2 NA, thereby avoiding a steric clash. The energy needed for such conformational changes could affect the affinity and neutralization of 1122A11 against recent NAs with Asn245 glycosylation. In the previous cryo-electron microscopy (cryo-EM) structure of FNI9 Fab in complex the N2 NA from A/Hong Kong/2671/2019 (H3N2) with Asn245 glycosylation, the N2 loop 242-252 and the Asn245 glycan appeared to be ordered but with dramatic conformational changes to avoid a steric clash between antibody and NA^20^, which is also the case in a recent cryo-EM structure of antibody DA03E17 in complex with N2 NA from A/Kansas/14/2017 (H3N2)^30^.

The primary focus of the functional analyses of 1122A11 here was to evaluate the dependence of engaging FcγRs for virus neutralization and protection, providing more mechanistic insight beyond the initial characterization of 1122A11 which demonstrated its potent prophylactic and therapeutic activity^21^. One potential shortcoming of this study was the use of mice to show *in vivo* protection following administration of the human mAb 1122A11 and 1122A11 LALAPG prophylactically and subsequent viral challenge. The pattern of expression of Fc-receptors on mouse cells and tissues does not necessarily mimic human expression^31^ and therefore, responses may be different than if a murine antibody were used. To overcome this shortcoming, perhaps using a mouse model that expresses human FcγRs^32^ would be more appropriate. Although mouse FcγRs and FcRn efficiently bind human IgG subclasses similarly to murine IgG subclasses^33^. Furthermore, the LALAPG mutation has been shown to greatly reduce mouse FcγRs engagement for murine antibodies^34^, as well as human IgG^35^ which would suggest that loss of Fc-FcR engagement is still valid in the mouse model used. The *in vivo* results showed that the LALAPG mutations in the Fc of 1122A11 did not result in differences in survival after viral challenge when prophylactically administered at 20 or 2 mg/kg. Future study of the impact LALAPG mutations and FcγR engagement requirement for the therapeutic activity of 1122A11 is warranted.

In the pursuit of creating a universal influenza vaccine, attention has been directed toward a few highly conserved epitopes on the virus. Notably, the HA stem region and the enzymatic active site of NA have been identified as promising targets. Presently, the efficacy of HA stem antibodies predominantly hinges on Fc-effector functions for *in vivo* activity^36, 37^. However, for NA active site antibodies, such as 1122A11 and FNI9^20^, binding to the NA active site enables direct anti-viral activity without requirement for Fc-effector functions to protect from lethal viral challenge. As a rare cross NA group protective mAb that was isolated from a licensed seasonal influenza vaccine, which is not optimized for the amount and quality of the NA, rather than from direct influenza virus infection, the structural and functional information from 1122A11 with N2 and other NAs provide insights for pan-NA antibody and vaccine design.

## Materials and Methods

### mAb production

A new antibody vector was created by cutting out the IgG1 constant region of AbVec hIgG1 (GenBank: FJ475055.1) using SalI and HindIII restriction enzymes and replacing it with a newly synthesized oligonucleotide (IDT) containing the L234A, L235A, and P329G (LALAPG) mutations. Insertion was confirmed via sequencing. The previously cloned mAb 1122A11^21^ underwent restriction digest with AgeI and SalI enzymes, was ligated into the new LALAPG vector, and was transformed into chemically competent DH5α E. coli cells. Resulting colonies were again sequenced to ensure the correct insertion. Purified plasmid DNA of either the parental heavy or the LALAPG heavy chain and parental light chains were transfected into 60–80% confluent HEK293T cells (ATCC) in 10 cm tissue culture treated dishes using JetPrime (Polyplus, Graffenstaden, France) as previously described^21^. Transfected cells were maintained for 8 days at 37°C in a 5% CO2 atmosphere while culturing in DMEM with 5% Fetal Clone II (Cytiva) and 1× antibiotic/antimycotic (Corning/Mediatech). Media were harvested and replaced every two to three days. Harvested media were concentrated using 100,000 MWCO Amicon Ultra centrifugal filters (Millipore-Sigma, Burlington, MA, USA) and then purified using Protein A magnetic beads (Promega, Madison, WI, USA) as described in the product literature. 1G01 and DA03E17 were synthesized based on previously reported VH and VL sequences (Genbank accession numbers: MN013068.1, MN013072.1, OP311730, OP311732). The F(ab’)2 of 1122A11 was generated by pepsin digestion of the intact antibody using the Pierce F(ab’)2 preparation kit as described in the product literature. Digestion was confirmed by mixing 2.5 µg of buffer exchanged digest with Laemmli buffer without β-mercaptoethanol (non-reducing conditions) and electrophoresis on a 10% TGX mini-Protein gel (Bio-Rad Laboratories) followed by staining with GelCode Blue Safe Protein Stain (Thermo Scientific). The stained gel can be found in Supplementary Fig. 4.

### Expression of Singapore16 N2 NA and 1122A11 Fab for crystal structure determination

The head domain of the N2 NA (residue 82-469) from A/Singapore/INFIMH-16-0019/2016 (H3N2) (Singapore16 N2, GISAID accession number EPI810155) was expressed in a baculovirus system, and purified as previously described^12^. Briefly, Singapore16 N2 NA head domain with A VASP tetramerization domain^38^ was cloned into a baculovirus transfer vector pFastbacHT-A (Invitrogen) with an N-terminal gp67 signal peptide, thrombin cleavage site, and His6-tag, and expressed in sf9 cells (Thermo Fisher Scientific). The secretion-expressed NA was purified from culture supernatant by metal affinity chromatography using Ni-nitrilotriacetic acid (Ni-NTA) resin (Qiagen). For crystal structure determination, the His_6_-tag and VASP tetramerization domain were cleaved from the N2 NA head domain by thrombin and the NA purified further by size exclusion chromatography in 20 mM Tris, pH 8.0, 150 mM NaCl and 0.02% NaN3.

The V_L_ and V_H_ regions of antibody 1122A11 Fab were cloned into the vector phCMV containing human lambda C_L_ region and human C_H_1 region appended to His_6_-tag, respectively^26^. The Fab was expressed by transiently co-transfection of the heavy and light chain vectors into ExpiCHO cells (Thermo Fisher Scientific) at 37°C with 8% CO_2_. The secretion-expressed Fab was purified from culture supernatant by metal affinity chromatography using Ni-NTA resin (Qiagen) and further purified by size exclusion chromatography using a Superdex 200 column (GE Healthcare) in 20 mM Tris, pH 8.0, 150 mM NaCl, and 0.02% NaN_3_.

### Crystal structure determination

The complex of 1122A11 Fab and Singapore16 N2 NA at 6.0 mg/mL was crystallized in 0.2 M tri-potassium citrate and 20% (w/v) polyethylene glycol 3350, and apo Singapore16 N2 NA at 9.9 mg/mL was crystallized in 0.1 M citric acid, pH 4.0, 2.4 M ammonium sulfate at 20 °C by the sitting drop vapor diffusion method using automated robotic crystal screening on our CrystalMation system (Rigaku). The crystals were flash-cooled at 100K in mother liquid and 15% ethylene glycol was added as cryoprotectant.

Diffraction data were collected at Stanford Synchrotron Radiation Lightsource (SSRL) beamlines 12-2 and 12-1 (Supplementary Table 1). Data were integrated and scaled with HKL3000^39^ Data and refinement statistics are summarized in Supplementary Table 1.

The crystal structure of 1122A11 Fab with Singapore16 N2 NA was determined by molecular replacement (MR) using the program Phaser^40^ using crystal structures of the Fab light and heavy chains (PDB codes 6B0S and 7K8Q) and the N2 NA from A/Tanzania/205/2010 (H3N2) (PDB code 4GZO) as input MR models. The refined Singapore16 N2 NA structure in its complex with 1122A11 Fab was used as the input MR model for structure determination of apo Singapore16 N2 NA. Initial rigid body refinement was performed in REFMAC5^41^ and further restrained refinement including TLS refinement was carried out in Phenix^9^. Model building between rounds of refinement was carried out with Coot^42^. Final statistics are summarized in Supplementary Table 1. The Asn245 glycosylation of apo Singapore16 N2 NA was built based on its well-defined electron density, but in its complex with 1122A11 Fab, there was only weak density for Asn245 glycosylation and not strong enough to be built at an atomic level. The quality of the structure was analyzed using JCSG validation suite (qc-check.usc.edu/QC/qc_check.pl) and MolProbity^43^. All structural figures were generated with PyMol (www.pymol.org).

### Expression of NA proteins for ELISA and MUNANA NA inhibition assay

For ELISA and MUNANA NA inhibition assays, the following NA proteins were expressed in a baculovirus system and purified as previously described^12^: The head domains of the N1 NA (residue 82-469) from A/Brevig Mission/1/1918 (H1N1) (GenBank accession number AF250356) and the N2 NA from A/Japan/305/1957 (H2N2) (GenBank accession number CY044327); the head domains plus stalk regions of the N2 NA from A/Hong Kong/1/1968 (H3N2) (GenBank accession number CY044263), the N2 NA from A/Singapore/INFIMH-16-0019/2016 (H3N2) (GISAID accession number EPI810155), the N3 NA from A/swine/Missouri/2124514/2006 (H2N3) (GenBank accession number EU258937), the N4 NA from A/mink/Sweden/E12665/84 (H10N4) (GenBank accession number AY207530), The N5 NA from A/duck/Alberta/60/1976 (H12N5) (GenBank accession number CY130080), The N6 NA from A/Hubei/29578/2016 (H5N6) (GenBank accession number CY044327), the N7 NA from A/Netherlands/219/2003 (H7N7) (GenBank accession number AY340079), the N8 NA from A/gyrfalcon/41088-6/2014 (H5N8) (GenBank accession number KP307986), the N9 NA from A/Hunan/02650/2016 (H7N9) (GISAID accession number EPI961189), the Victoria lineage influenza B NA from B/Colorado/06/2017 (GISAID accession number EPI969379), and the Yamagata lineage influenza B NA from B/Phuket/3073/2013 (GISAID accession number EPI529344). Briefly, the NA constructs with A VASP tetramerization domain^38^ was cloned into a baculovirus transfer vector pFastbacHT-A (Invitrogen) with an N-terminal gp67 signal peptide, thrombin cleavage site, and His_6_-tag, and expressed in sf9 cells. The secretion-expressed NAs were purified from culture supernatant by metal affinity chromatography using Ni-nitrilotriacetic acid (Ni-NTA) resin (Qiagen) and buffered in 20 mM Tris, pH 8.0, 150 mM NaCl and 0.02% NaN_3_.

### Enzyme-linked immunosorbent assay (ELISA)

ELISA was performed as previously described^21^ and modified such that the coating buffer and dilution buffer was PBS or DPBS with calcium and magnesium (Corning Mediatech). ELISA plates (Nunc MaxiSorp; Thermo Fisher Scientific, Grand Island, NY, USA) were coated with recombinant NA at 2 µg/mL. mAbs were diluted in PBS or DPBS, and binding was detected with HRP-conjugated anti-human IgG (Jackson ImmunoResearch, West Grove, PA, USA).

### FcγRI binding

Both 1122A11 and 1122A11–LALAPG were both coated onto ELISA plates (Nunc MaxiSorp; Thermo Fisher Scientific, Grand Island, NY, USA) at 1 µg/mL. Plates were sealed and stored at 4°C overnight before being washed 3× in PBS. Biotinylated Human Fc gamma RI/CD64 (Acro Biosystems, Newark, DE, USA) was diluted to 1, 0.1, and 0 µg/mL, and 50 µL was added to wells in triplicate for each antibody concentration. After 1 h of shaking at room temperature, plates were washed 5× in PBS with 0.05% Tween 20, and 50 µL of 2 µg/mL Streptavidin–horseradish peroxidase (Jackson ImmunoResearch, West Grove, PA, USA) was added and incubated for 1 h, with shaking at room temperature. Plates were again washed 5× in PBS with 0.05% Tween 20, and 50 µL of TMB substrate (SureBlue; KPL, SeraCare, Milford, MA) was added to wells and the plates incubated for 5 min at room temperature. To stop the reaction, 50 µL of 1N HCl was added to wells and absorbance was read at 450 nm.

### Enzyme linked lectin assay (ELLA)

ELLA were performed essentially as previously described by Bernard et al.^44^. In short, 96-well flat bottom immune plates (Nunc, ThermoFisher) were coated with 50 µg/mL Fetuin in PBS, sealed, and refrigerated overnight at 4°C. The next day, plates were washed three times with PBS–Tween (0.05%). Assay optimization was performed using rNA of A/Hong Kong/2286/2017 (BEI Resources, NR-52032) by conducting 1:2 serial dilutions in DPBS (with calcium and magnesium) with 1% BSA to cover the range of 0.003906 to 8 µg/mL. Plates were sealed and incubated for 18 h at 37°C before being washed 6 times with PBS–Tween. A total of 50 µL of peroxidase-conjugated lectin of *Arachis hypogaea* (peanut) was added to plates (2 µg/mL) and incubated for 2 h in the dark. Plates were washed three times, and 50 µL of TMB substrate (SureBlue; KPL, SeraCare) was added to wells and plates incubated for 20 min at room temperature. To stop the reaction, 50 µL of 1N HCl was added to wells and absorbance was read at 450 nm. A concentration of rNA was selected that was approximately 90% of the peak of the linear range of the resulting curve (∼0.03 µg/mL) for inhibition assays. Both the 1122A11 parental and LALAPG variant mAbs were diluted 1:2 on a washed fetuin-coated plate to cover a range from 0.006 to 12 µg/mL. The rNA was then added at the predetermined concentration to all but the “background” wells. Plates were sealed and incubated for approximately 20 h at 37°C before being washed and developed as described above. Optical densities were corrected using the mean of the “background” wells, and then percent inhibition was calculated from the mean of wells incubated with the rNA alone. The IC_50_ was determined using non-linear regression in GraphPad Prism (Version 9.5.1).

### MUNANA NA inhibition assay

Neuraminidase inhibition assays were carried out similar to the procedure described previously^45^ except recombinant NA were used instead of influenza virus. Because NA activity is influenced by both calcium and the presence of protein, 3% (wt./volume) bovine serum albumin was added to the 1x assay buffer (33.3 mM 2-(N-morpholino) ethanesulfonic acid (MES) and 4 mM CaCl_2_, pH 6.5) so that inhibition by mAbs could be assessed without further activation of the enzyme. To determine optimal activity of each NA, each recombinant protein was serially diluted 1:1 in 1x assay buffer with an initial concentration of 10 µg/mL before adding 50 µl of 300 µM 2′-(4-Methylumbelliferyl)-α-D-*N*-acetylneuraminic acid (MUNANA) (Sigma, St. Louis, MO) to each well and incubating for 1 h at 37⁰C. Optimal NA concentration was determined to be 80% of the highest reading in the linear range (Supplementary Table 5) of the fluorescence after 355 nm excitation and 460 nm emission on a Cytation3 (BioTek, Winooski, VT). The optimal concentration was then used in subsequent inhibition experiments with serial 1:1 dilutions of 1122A11 and 1G01 (starting concentration either 10 or 2 µg/mL). Mean fluorescence of MUNANA only wells was subtracted as background from all remaining wells and then the mean of NA + MUNANA was used as “100% activity.” Percent activity was then calculated for each well with dilutions of antibodies. Non-linear regression of percent activity versus antibody concentration was used to calculate the 50% of the inhibitory concentration (IC_50_) using GraphPad Prism (Ver. 10). When inhibition was not detected, the IC_50_ is reported as >10 µg/mL and when inhibition was obviously less than the lowest concentration of antibody, the IC_50_ was expressed as less than (<) that concentration.

### Antibody-dependent cellular phagocytosis (ADCP)

ADCP activity of the hMAbs was measured as previously described^46, 47^ with slight modifications. Briefly, NA of A/Singapore/INFIMH-16-0019/2016 was biotinylated with the Biotin-XX Microscale Protein Labeling Kit (Life Technologies, Carlsbad, CA, USA). A total of 0.25 μg of biotinylated Ag or ∼0.32 μg of BSA (used as a baseline control in an equivalent number of Ag molecules/bead) was incubated overnight at 4 °C with 1.9 × 10^6^ yellow-green neutravidin-fluorescent beads (Life Technologies) per reaction in a 25 μL of final volume. Antigen-coated beads were subsequently washed twice in PBS–BSA (0.1%) and transferred to a 5 mL Falcon round bottom tube (Thermo Fisher Scientific, Waltham, MA, USA). hMAbs, diluted at 0.5 or 5 μg/mL, were added to each tube in a 20 μL of reaction volume and incubated for 2 h at 37 °C in order to allow Ag–Ab binding. Then, 250,000 THP-1 cells (human monocytic cell line obtained from NIH AIDS Reagent Program) were added to the cells and incubated for 3 h at 37 °C. At the end of incubation, 100 μL 4% paraformaldehyde was added to fix the samples. Cells were then assayed for fluorescent bead uptake by flow cytometry using a BD Biosciences Symphony. The phagocytic score of each sample was calculated by multiplying the percentage of bead positive cells (frequency) by the degree of phagocytosis measured as mean fluorescence intensity (MFI) and divided by 10^6^. Values were normalized to background values (cells and beads without mAb) and an isotype control to ensure consistency in values obtained on different assays. Finally, the phagocytic score of the testing hMAb was expressed as the fold increase over BSA-coated beads.

### Microneutralization assay

The neutralizing activity of both 1122A11 and 1122A11–LALAPG mAbs against A/Uruguay/716/07 (H3N2) virus (BEI Resources) was performed using a virus neutralization assay as previously described^21^. Briefly, confluent monolayers of MDCK cells (ATCC (5 × 10^4^ cells/well, 96-well plate format, quadruplicates)) were infected with 200 plaque forming units (PFU)/well of the indicated virus. After 1 h of viral adsorption, cells were maintained in a post-infection medium (DMEM, 0.3% BSA, 1% PSG, and 1 μg/mL tosylsulfonyl phenylalanyl chloromethyl ketone (TPCK)-treated trypsin (Sigma) containing two-fold serial dilutions (starting concentration of 50 µg/mL) of the mAbs and incubated at 37°C with 5% CO2. At 72 h p.i., virus neutralization titer (NT) was evaluated following crystal violet staining of the cells and expressed as the lowest concentration of the mAb to prevent the virus-induced cytopathic effect (CPE).

### In vivo study

The prophylactic activities of both 1122A11 and 1122A11–LALAPG mAbs were tested against X31 in mice. Briefly, female C57BL/6 mice (7 weeks of age) were purchased from the Jackson Laboratory and maintained in the animal facility at Texas Biomedical Research Institute under specific, pathogen-free conditions and ABSL2 containment. Mice were intraperitoneal treated with a single dose of 2 or 20 mg/kg of each mAb, isotype control, or PBS (mock treated) 24 h prior to viral challenge. For virus infection, mice were anesthetized following gaseous sedation in an isoflurane chamber and inoculated intranasally with 10X MLD_50_ (3×10^4^ PFU)/mouse of X31 (A recombinant influenza A/Puerto Rico/8/1934 (H1N1) containing the HA and NA segments of A/Hong Kong/1/1968 (H3N2) virus was obtained from Dr. David Topham and grown in MDCK cells as previously described^48^). At days 2 and 4 p.i., viral replication was determined in the lungs of the infected mice. Briefly, four mice from each experimental group were euthanized by administration of a lethal dose of avertin and exsanguination, and lungs were surgically extracted and homogenized in 1 mL of PBS using Precellys tissue homogenizer (Bertin Instruments, Rockville, MD, USA). Virus titers (PFU/mL) were determined by standard plaque assay as previously described^21^. Geometric mean titers and data representation were performed using GraphPad Prism (v9.0). For the body weight and survival study, mice (N = 5, group) were monitored daily for 10 days p.i. for morbidity (body weight) and mortality (survival rate). Mice showing a loss of more than 25% of their initial body weight were defined as reaching the experimental endpoint and humanely euthanized. All animal protocols were approved by the Texas Biomedical Research Institute (IACUC 1785 MU 0).

### Statistics and Reproducibility

Means and standard deviations of replicate measures are presented in figures and tables. One-way analysis of variance (ANOVA) and Tukey’s test for multiple-comparison correction was used to determine the statistical significance of the *in vivo* viral titers in Graphpad Prism Version 9.0. For non-detection, 20 FFU (the limit of detection) was assigned to those values for statistical analysis based on the initial dilution of lung homogenate (1:10). This results in a bias of the calculated mean, such that the value is probably higher than what could potentially be measured. Significant differences between resulting means was declared at *P* < 0.05. No data was excluded from the analysis. Animal sample size of four animals per group per timepoint was based on previously published work^21, 49^. Animals were randomly allocated to each group to ensure reproducibility.

## Author Contributions

L.M.-S., I.A.W. and J.J.K. conceived the project. X.Z., A.M.K., M.S.P., and W.Y. performed laboratory investigations. X.Z., A.M.K., M.S.P., L.M.-S., I.A.W., and J.J.K. analyzed data and visualized results. X.Z., J.J.K., and I.A.W. wrote the original draft. All authors revised and approved the final manuscript. All authors have read and agreed to the submitted version of the manuscript.

## Institutional Review Board Statement

Studies with influenza virus were approved by the Texas Biomedical Research biosafety (BSC21-017) and recombinant DNA (RDC21-017) committees. Animal studies were approved by the Texas Biomedical Research Institute’s Institutional Animal Care and Use Committee (IACUC 1785 MU 0).

## Data Availability Statement

The authors confirm that the data supporting the findings of this study are available within the article and supplemental material. The atomic coordinates and structure factors are being deposited in the Protein Data Bank (PDB) under accession codes 9MQW and 9MQV for apo Singapore16 N2 NA and its complex with 1122A11 Fab.

## Acknowledgments

We would like to thank the Biodefense and Emerging Infections Research Resources Repository (BEI Resources) and the International Reagent Resource (IRR) for providing reagents used in this study. We thank Henry Tien for automated robotic crystal screening. The SSRL is a Directorate of SLAC National Accelerator Laboratory and an Office of Science User Facility operated for the U.S. Department of Energy Office of Science by Stanford University. The SSRL Structural Molecular Biology Program is supported by the DOE Office of Biological and Environmental Research, and by the National Institutes of Health, National Institute of General Medical Sciences (including P41GM103393) and the National Center for Research Resources (P41RR001209). This research was funded in-part by the National Institutes of Health, National Institute of Institute of Allergy and Infectious Diseases (5R01AI145332). L.M.-S. research on influenza is supported by a grant from the American Lung Association (ALA). A.M.K. was partially funded by a Texas Biomedical Research Institute Forum award.

