## Supplemental Information for "Structure and function of a cross-neutralizing influenza neuraminidase antibody that accommodates recent N2 NA Asn245 glycosylation"

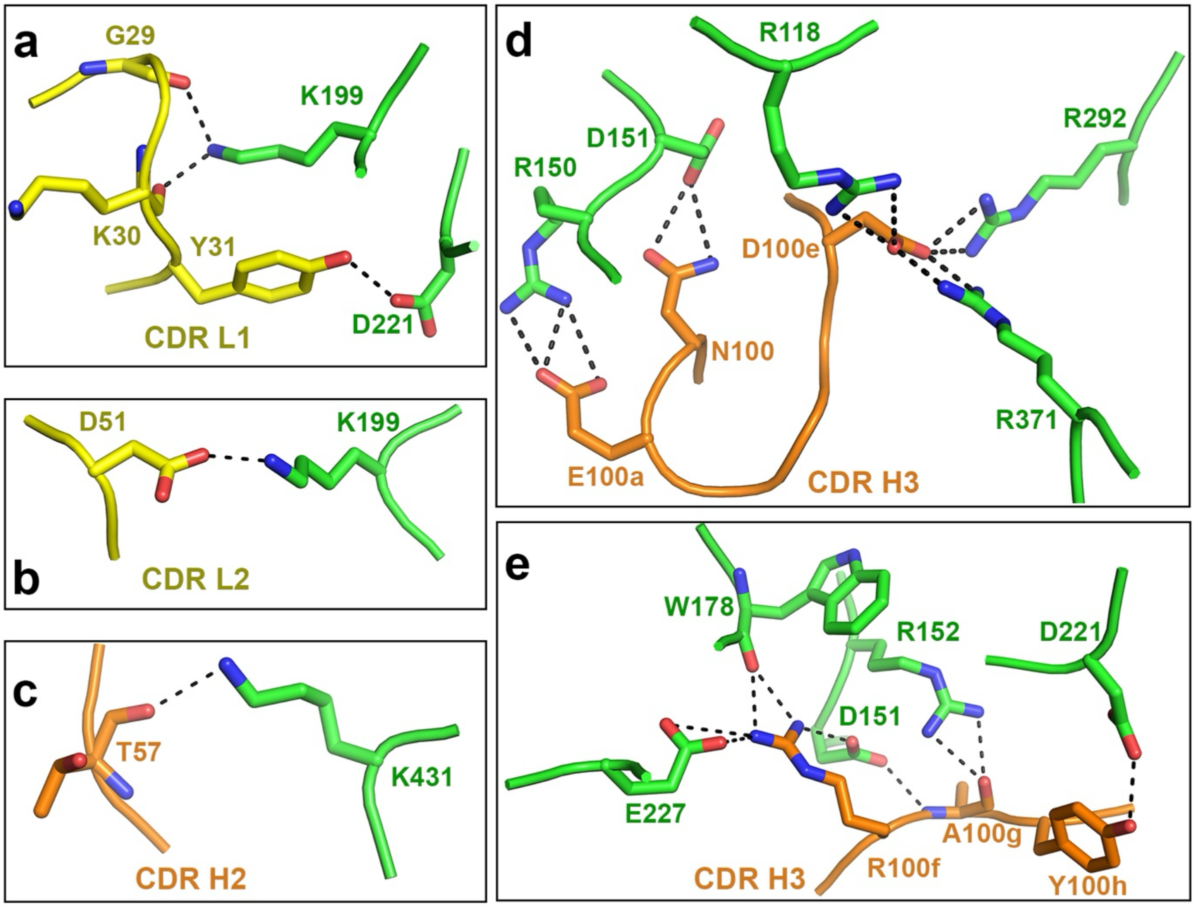


**Supplementary Fig. 1. Hydrogen bond and salt-bridge interactions of 1122A11 Fab with Singapore16 N2 NA from the crystal structure at 2.2 Å resolution. a** The antibody CDR L1 interaction with the NA. **b** CDR L2 interaction with the NA. **c** CDR H2 interactions with the NA. **d** and **e** CDR H3 interaction with the NA. The NA residues are colored in green, and the Fab light and heavy chains are shown in yellow and orange, respectively.

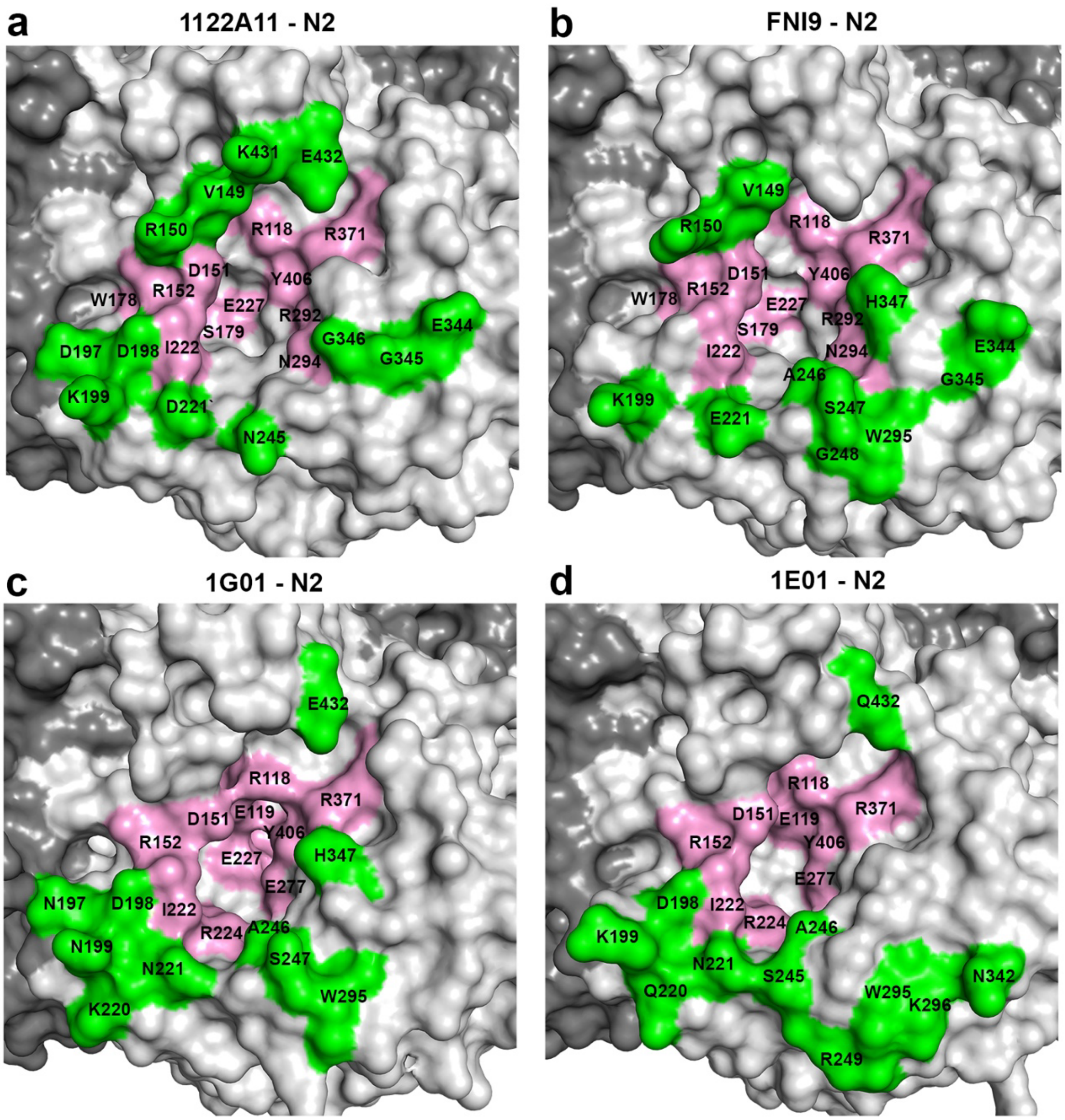


**Supplementary Fig. 2. Epitope comparison of 1122A11 and other human mAbs with N2 NAs. a** The epitope of 1122A11 is mapped onto Singapore16 N2 NA (H3N2) (X-ray, 2.2 Å resolution). **b** The epitope of FNI9 is mapped onto N2 NA from A/Tanzania/205/2010 (H3N2) (cryo-EM, 2.9 Å resolution, PDB code 8G3N). **c** The epitope of 1G01 is mapped onto N2 NA from A/Indiana/10/2011 (H3N2v) (cryo-EM, 3.0 Å resolution, PDB code 8GAT). **d** The epitope of 1E01 is mapped onto N2 NA from A/Japan/306/1957 (H2N2) (X-ray, 2.45 Å resolution, PDB code 6Q20). One NA protomer is colored in light grey, and other NA protomers are in dark grey. The molecular surface depicting the epitope residues colored in green with epitope residues colored in pink for conserved resides in all subtypes of influenza A and B NAs.

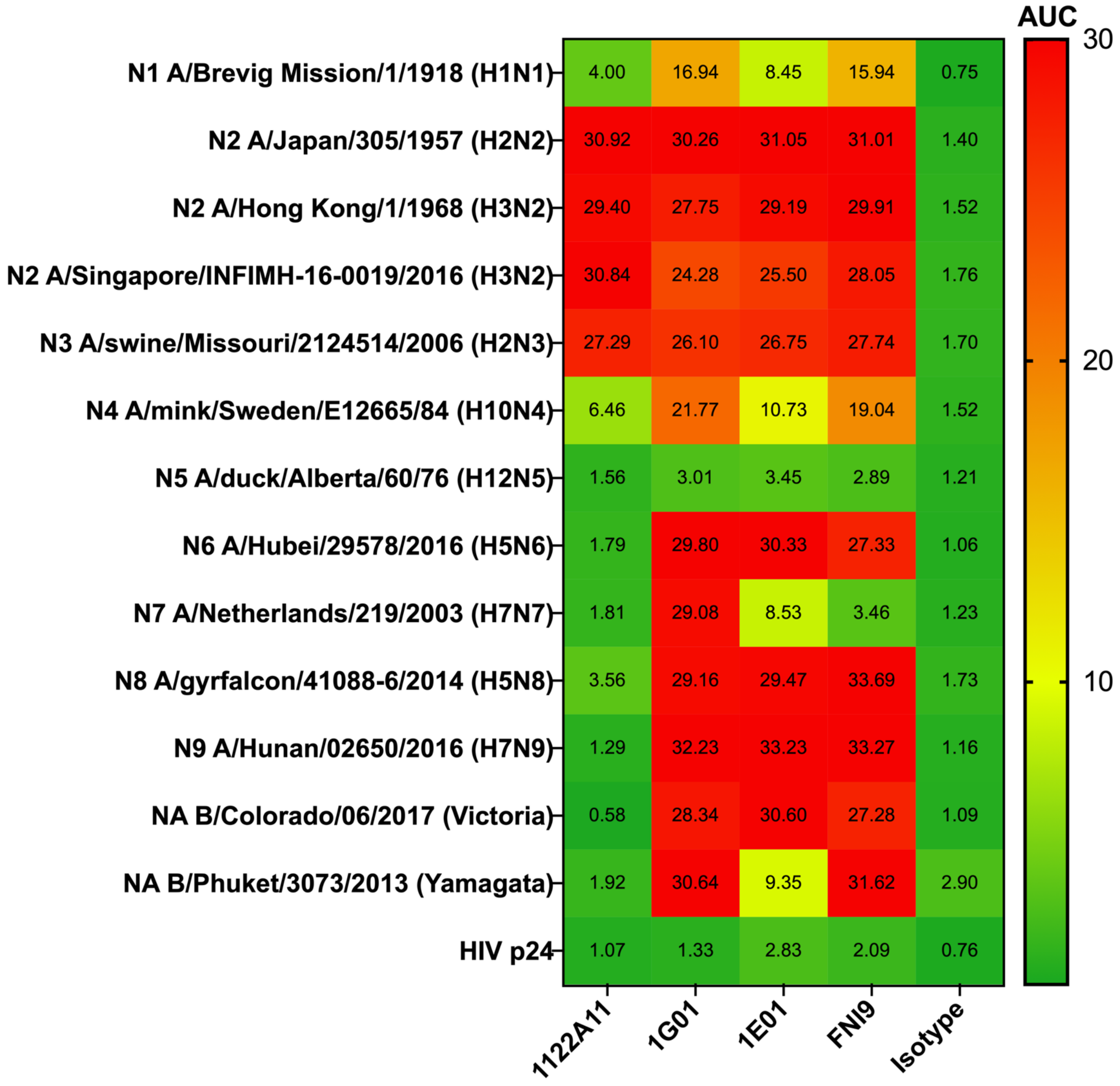


**Supplementary Fig. 3. Breadth of mAb 1122A11 binding to recombinant NA when no calcium is present in coating and dilution buffers.** Heat map of 1122A11 binding presented as area under the curve (AUC) with red representing highest binding and green lowest binding across all data) to recombinant NA proteins in ELISA with positive control pan-NA antibodies 1G01, 1E01 and FNI9, as well as a negative control antibody Isotype.

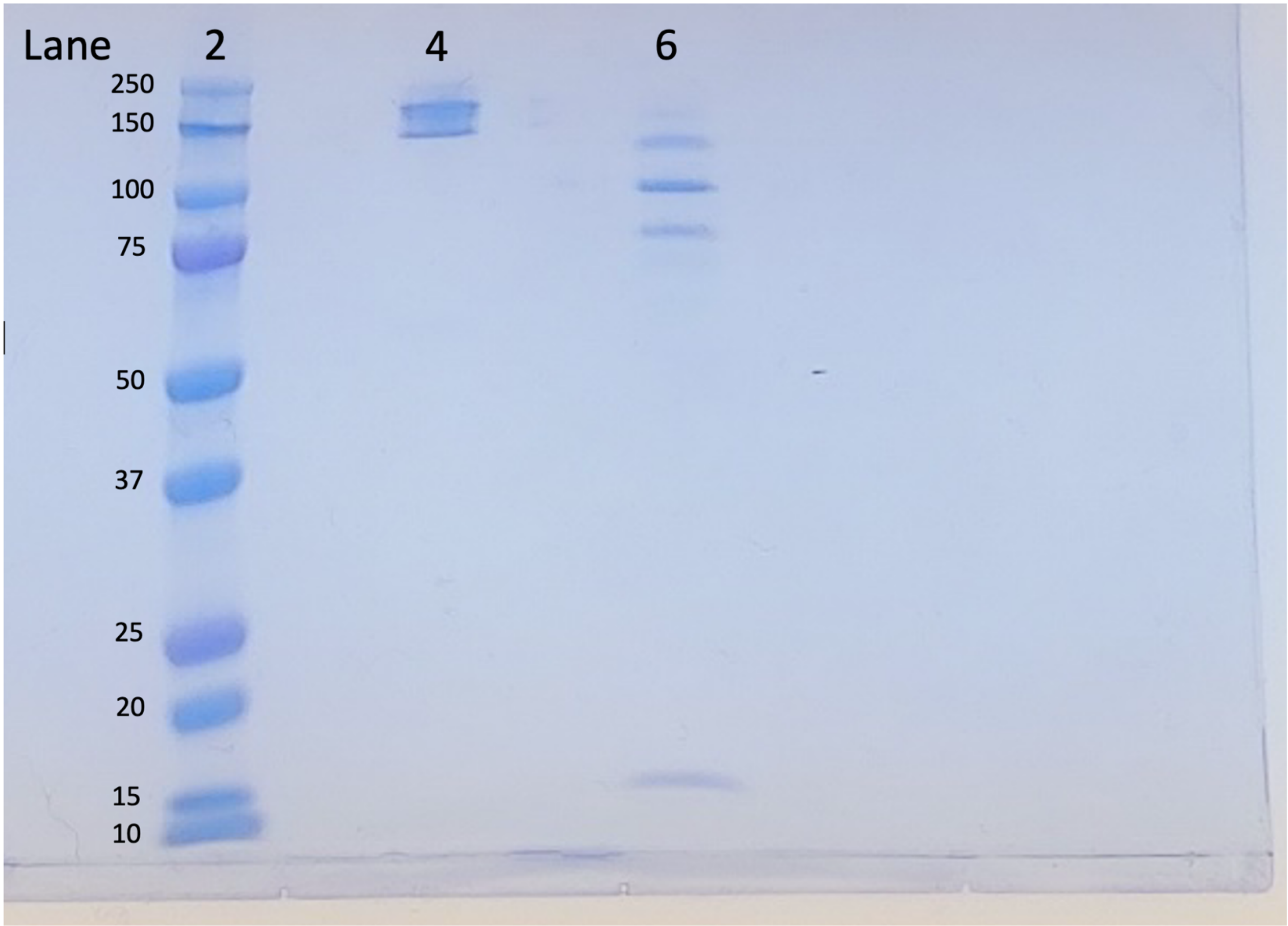


**Supplementary Fig. 4. Gel comparing intact mAb 1122A11 with the resulting F(ab')2 after pepsin digestion.** Lane 2 contains Precision Plus Protein Standard (Bio-Rad). 2.5 μg of protein was loaded for 1122A11 (lane 4) and 1122A11 F(ab')2 (lane 6) under non-reducing conditions. Following electrophoresis, gel was washed 3 x 5 minutes in water and then stained for 45 minutes with GelCode Blue Safe Protein Stain (Pierce). Gel was destained for 3 hours with 3 water changes.

### Supplementary Table 1. Data collection and refinement statistics

| Data set | **Singapore16 N2 NA + 1122A11 Fab** | **Singapore16 N2 NA** |
|---|---|---|
| **Data Collection** | | |
| X-ray source | SSRL 12-2 | SSRL 12-1 |
| Wavelength (Å) | 0.97946 | 0.97946 |
| Space group | P4 | I422 |
| Unit cell (Å) | *a* = *b* = 115.9, *c* = 76.8 | *a* = *b* = 137.3, *c* = 155.5 |
| angle (º) | 90, 90, 90 | 90, 90, 90 |
| Resolution (Å)[a] | 38.63-2.20 (2.25-2.20) | 48.55-2.30 (2.34-2.30) |
| Unique reflections[a] | 51,700 (3,381) | 32,018 (1,258) |
| Redundancy[a] | 12.0 (6.2) | 17.6 (5.1) |
| Average $I/\sigma(I)$[a] | 18.1 (1.1) | 12.6 (0.7) |
| Completeness (%)[a] | 99.6 (98.3) | 95.4 (76.7) |
| $R_{sym}$[a,b] | 0.13 (>1.0) | 0.26 (>1.0) |
| $R_{pim}$[a,b] | 0.04 (0.47) | 0.06 (0.63) |
| $CC_{1/2}$[a] | 0.999 (0.536) | 0.993 (0.629) |
| No. molecules per ASU[c] | 1 | 1 |
| **Refinement** | | |
| Resolution (Å)[a] | 38.63-2.20 (2.24-2.20) | 48.55-2.30 (2.36-2.30) |
| Reflections in refinement[a] | 51,678 (2,626) | 31,802 (2,008) |
| Refined residues | 818 | 387 |
| Refined waters | 365 | 163 |
| $R_{cryst}$[a,d] | 0.169 (0.284) | 0.175 (0.339) |
| $R_{free}$[a,e] | 0.204 (0.338) | 0.214 (0.346) |
| *B*-values (Å$^2$) | | |
| Protein | 50 | 47 |
| NA | 37 | 47 |
| Fab variable | 47 | - |
| Fab constant | 77 | - |
| Water | 47 | 52 |
| Wilson *B*-values (Å$^2$) | 39 | 41 |
| Ramachandran values (%)[f] | 97.4, 0.3 | 96.1, 0 |
| r.m.s.d. bond (Å) | 0.003 | 0.005 |
| r.m.s.d. angle (deg.) | 0.59 | 0.83 |
| PDB codes | 9MQV | 9MQW |

[a] Parentheses denote outer-shell statistics.

[b] $R_{sym} = \sum_{hkl}\sum_i |I_{hkl,i} - \langle I_{hkl}\rangle| / \sum_{hkl}\sum_i I_{hkl,i}$ and $R_{pim} = \sum_{hkl}[1/(N-1)]^{1/2}\sum_i |I_{hkl,i} - \langle I_{hkl}\rangle| / \sum_{hkl}\sum_i I_{hkl,i}$, where $I_{hkl,i}$ is the scaled intensity of the i$^{th}$ measurement of reflection *h*, *k*, *l*, $\langle I_{hkl}\rangle$ is the average intensity for that reflection, and *N* is the redundancy. $R_{pim} = \Sigma_{hkl} (1/(n-1))^{1/2} \Sigma_i | I_{hkl,i} - \langle I_{hkl}\rangle | / \Sigma_{hkl} \Sigma_i I_{hkl,i}$, where n is the redundancy

[c] No. molecules for complexes refers to number of NA protomers plus antibody Fab or number of NA protomers per asymmetric unit (ASU).

[d] $R_{cryst} = \sum_{hkl} |F_o - F_c| / \sum_{hkl} |F_o|$, where $F_o$ and $F_c$ are the observed and calculated structure factors.

[e] $R_{free}$ was calculated as for $R_{cryst}$, but on 5% of data excluded before refinement.

[f] The values are percentage of residues in the favored and outliers regions analyzed by MolProbity[1].

**Supplementary Table 2. Sequence alignment around 1122A11 epitope residues in Singapore16 N2 NA for NA strains tested binding to 1122A11 and comparison to the N2 NA epitopes of FNI9, 1G01 and 1E01.**

| | 118 | 149 | 178 | 197 | 221 227 | 245 |
|---|---|---|---|---|---|---|
| A/Singapore/INFIMH-16-0019/2016(H3N2) | VTREP | NTVRDRTP | IAWSSS | TGDDKNA | SKDILRTQESE | DGNATG |
| A/Brevig Mission/1/1918(H1N1) | .I... | G..K..S. | V...A. | S.P.NG. | RNN........ | ..PSN. |
| A/Japan/305/1957(H2N2) | ..... | G.IH..I. | V..... | ....R.. | .QN........ | ..S.S. |
| A/Hong Kong/1/1968(H3N2) | ..... | D.IH..I. | ...... | ....... | .QN........ | ..S.S. |
| A/Uruguay/716/2007(H3N2) | ..... | D....... | ...... | ....... | ..E........ | ..S.S. |
| A/Hong Kong/2286/2017(H3N2) | ..... | ........ | ...... | ....... | .N......... | ...... |
| A/swine/Missouri/2124514/2006(H2N3) | ..... | G.IK.... | ...... | ..N.ND. | R.......... | ..P.AN |
| A/gyrfalcon/Washington/41088-6/2014(H5N8) | .I... | G..K..S. | V...AT | ..P.SK. | AG........S | ..P.NR |
| A/mink/Sweden/E12665/84(H10N4) | .I... | G..K..S. | V...AT | S.P.AT. | RGN.M...... | ..PSDA |
| A/duck/Alberta/60/1976(H12N5) | .I... | ...K..S. | V...AT | S.A.DD. | R.Q........ | ..P.NS |
| A/Hubei/29578/2016(H5N6) | ..... | G.IH..G. | .G...T | S.PNN.. | VGN.......S | ..P.NN |
| A/Netherlands/219/2003(H7N7) | ..... | G.IH...A | VG...T | Q.NND.. | ARN........ | ..S.SS |
| A/Hunan/02650/2016(H7N9) | ..... | G.IH..SQ | .G...T | S.PNN.. | ARN........ | ..P... |
| B/Colorado/06/2017(Victoria) | II... | G.RG..NK | A...G. | D.P.N.. | ANN.......A | ..S.S. |
| B/Phuket/3073/2013(Yamagata) | II... | G.RE..NK | A...G. | D.P.S.. | A.N.......A | ..P.S. |
| A/Indiana/10/2011(H3N2)- 1G01 | ..... | ...H.... | M..... | ..N.N.. | ..N.......S | ..S.S. |
| A/Japan/305/1957(H2N2)- 1E01 | ..... | G.IH..I. | V..... | ....R.. | .QN........ | ..S.S. |
| A/Tanzania/205/2010(H3N2)- FNI9 | ..... | ........ | ...... | ....... | ..E.......S | ..S.S. |

| | 277 | 292 | 344 | 371 | 406 | 431 |
|---|---|---|---|---|---|---|
| A/Singapore/INFIMH-16-0019/2016(H3N2) | EEC | VCRDNWK | NE--EGG-HG | TSRLG | SGYSG | GR-KEET |
| A/Brevig Mission/1/1918(H1N1) | ... | ......H | ----N.A-N. | S..S. | ..... | .Q-PK.N |
| A/Japan/305/1957(H2N2) | ... | I...... | ..--R.N-P. | D..S. | ..... | ..-PQ.. |
| A/Hong Kong/1/1968(H3N2) | ... | I...... | ..--R.N-Q. | DL.S. | ..... | ..-.Q.. |
| A/Uruguay/716/2007(H3N2) | ... | ....... | ..--...-.. | ..... | ..... | ..-.... |
| A/Hong Kong/2286/2017(H3N2) | ... | ....... | ..--...-.. | ..... | ..... | ..-.... |
| A/swine/Missouri/2124514/2006(H2N3) | ... | I...... | .V--N..-P. | SG.S. | ..... | ..-PNKN |
| A/gyrfalcon/Washington/41088-6/2014(H5N8) | ... | ......T | ----NQ.-Y. | ...S. | ..... | ..-P..R |
| A/mink/Sweden/E12665/84(H10N4) | ... | ......R | ----K.R-Y. | E..S. | ..... | .Q-PK.K |
| A/duck/Alberta/60/1976(H12N5) | ... | ......N | -GGSGTNNY. | S..S. | ..... | .KPE.R. |
| A/Hubei/29578/2016(H5N6) | ... | ....... | -I--G.S-PD | D..S. | ..... | ..P..ID |
| A/Netherlands/219/2003(H7N7) | ... | ......Q | -T--GSP-GV | R..S. | ..... | ..-P..A |
| A/Hunan/02650/2016(H7N9) | ... | T.....Q | PG--N-NNN- | A..S. | ..... | ..P..DK |
| B/Colorado/06/2017(Victoria) | ... | A....RY | GD--K.S-G. | .E.M. | GW..F | DG----G |
| B/Phuket/3073/2013(Yamagata) | ... | A....SY | GD--E.S-G. | .K.M. | GW..F | DG----G |
| A/Indiana/10/2011(H3N2)- 1G01 | ... | ....... | ..--K..-.. | .L... | ..... | ..-.... |
| A/Japan/305/1957(H2N2)- 1E01 | ... | I...... | ..--R.N-P. | D..S. | ..... | ..-PQ.. |
| A/Tanzania/205/2010(H3N2)- FNI9 | ... | ....... | ..--...-.. | ..... | ..... | ..-.... |

Antibody 1122A11 contact residues in Singapore16 N2 NA are highlighted in green. In ELISA binding experiments, the tested influenza A strain N1, N2, N3, N8 NAs on the top part were reactive to mAb 1122A11, but not or minimal to the tested influenza A strain N4, N5, N6, N7 and N9 as well as two flu B NAs in the middle part of this table. For comparison, the contact residues in the N2 NAs from A/Tanzania/205/2010 (H3N2), A/Indiana/10/2011 (H3N2) and A/Japan/305/1957 (H2N2) in their complex structures with corresponding antibodies FNI9 (PDB code 8G3N), 1G01 (PDB code 8GAT) and 1E01 (PDB code 6Q20) are also highlighted in magenta, cyan and swamp green, respectively. "." denotes the same residue as Singapore16 N2 NA, "-" denotes sequence deletion as aligned to Singapore16 N2 NA.

| A/Uraguay/716/07 (H3N2) | |
|---|---|
| mAb | $NT_{50}$ (µg/ml) |
| 1122A11 | 1.56 |
| 1122A11-LALAPG | 0.78 |
| 1G01 | 6.25 |
| DA03E17 | 6.25 |

**Supplementary Table 3. Ability of 1122A11-LALAPG to inhibit viral infection.** Microneutralization assay. MDCK cells were infected with A/Uruguay/716/07 (H3N2) virus and then incubated with two-fold serial dilutions (starting concentration, 50 µg/mL) of the mAbs. Virus NT was evaluated 72 h post infection following crystal violet staining and expressed as the lowest concentration of the mAb to prevent virus-induced CPE. Mock-infected cells and viruses in the absence of mAb were used as internal controls.

| A/Uraguay/716/07 (H3N2) | |
|---|---|
| mAb | $NT_{50}$ (µg/ml) |
| 1122A11 | 0.29 |
| 1122A11-LALAPG | 0.22 |
| 1122A11 – $F(ab')_2$ | 0.16 |

**Supplementary Table 4. Ability of 1122A11-LALAPG to inhibit viral infection.** Microneutralization assay. MDCK cells were infected with A/Uruguay/716/07 (H3N2) virus and then incubated with two-fold serial dilutions (starting concentration, 50 µg/mL) of the mAbs. Virus NT was evaluated 72 h post infection following crystal violet staining and expressed as the lowest concentration of the mAb to prevent virus-induced CPE. Plates were rinsed with water and then methanol was added. After 20 minutes incubation at RT, the optical density at 560 nm was measured using a GloMax Discover system. The $NT_{50}$ was cucullated by GraphPad Prism. Mock-infected cells and viruses in the absence of mAb were used as internal controls.

| Recombinant NA | Concentration (μg/ml) to reach 80% of maximum relative fluorescence |
|---|---|
| A/Brevig Mission/1/1918 (H1N1) | 0.045 |
| A/Japan/305/1957 (H2N2) | 0.008 |
| A/Hong Kong/1/1968 (H3N2) | 0.044 |
| A/Singapore/INFIMH-16-0019/2016 (H3N2) | 0.021 |
| A/Hong Kong/2286/2017 (H3N2) | 0.055 |
| A/swine/Missouri/2124514/2006 (H2N3) | 0.087 |
| A/mink/Sweden/E12665/84 (H10N4) | 0.169 |
| A/duck/Alberta/60/76 (H12N5) | 1.272 |
| A/Hubei/29578/2016 (H5N6) | 0.033 |
| A/Netherlands/219/2003 (H7N7) | 0.033 |
| A/gyrfalcon/41088-6/2014 (H5N8) | 0.011 |
| A/Hunan/02650/2016 (H7N9) | 0.044 |
| B/Colorado/06/2017 (Victoria) | 0.055 |
| B/Phuket/3073/2013 (Yamagata) | 0.035 |

**Supplemental Table 5. Determining ideal concentration of neuraminidase to be used in the MUNANA assay.** Maximum fluorescence was determined by initial serial dilution of 4-methylumbelliferone in MUNANA assay buffer, incubating with 300 μM 4-Methylumbelliferyl-N-acetyl-α-D-neuraminic acid for 30 minutes at 37ºC, then adding 100 μl Basic Stop solution and taking a reading in the Cytation3 (BioTek) with excitation wavelength of 355 nm and emission wavelength of 460 nm. Data points from high concentrations were removed stepwise until a linear regression could be determined with a *P* value for fit of > 0.95. Eighty percent of the highest relative fluorescence (for our Cytation3 and RFU of 19810 was used) for the linear range was then used as a target for determining optimum recombinant neuraminidase activity.
